# Apparent stabilizing selection explains micro and macroevolution of *Drosophila* wings

**DOI:** 10.64898/2026.01.08.698402

**Authors:** Anneli Brändén, Stephen P. De Lisle

## Abstract

Evolution over deep time is often slow relative to neutral or directional expectations. A striking example is provided by *Drosophila* wing shape, which shows substantial genetic variance yet comparatively little macroevolutionary change. Using a large-scale fitness assay, we find pervasive stabilizing selection across dimensions of *D. melanogaster* wing shape that is proportional to standing variation, driven by variance in male fitness, and likely generated by pleiotropy. Parameterizing a macroevolutionary Ornstein-Uhlenbeck model using our selection estimate recapitulates observed among species variance under reasonable effective population sizes, suggesting a role for apparent stabilizing selection in constraining wing shape evolution over deep time.

## Main Text

Stabilizing selection is a fundamental process in any explanation of evolutionary change over long timescales. In most lineages, ample genetic variance means that expectations for evolution under neutral genetic drift or directional selection result in wildly-high predictions for divergence over even moderate timescales^1–3^. Without discounting a supporting role for developmental and genetic constraints, no explanation for macroevolutionary stasis is complete without invoking a leading role for some form of stabilizing selection^3–5^. Surprisingly, there have been remarkably few^6,7^ empirical attempts to directly assess the role of stabilizing selection in generating patterns of stasis.

Here we report the results of a large-scale assay (phenotypic scores on 91,632 offspring from 2110 parents) of individual fitness and wing phenotype in a lab-adapted lineage of fruit fly, *Drosophila melanogaster*. Fruit fly wings (Figure 1) exemplify how the perennial puzzle of macroevolutionary stasis has itself evolved when confronted with more (and more sophisticated) datasets. Wing shape has remained remarkably unchanged across the Drosophilidae^4^ and beyond^8,9^, despite substantial genetic variance across all dimensions of measured wing shape^10^. Intriguingly, within-species patterns of mutational and standing genetic variance are conserved in the form of among-species evolutionary rate; wing shape combinations with high mutational input exhibit high standing genetic variance within species and high macroevolutionary rate among species^11^. These studies highlight some role for mutational constraints^11^ and developmental bias^8,9^ in linking micro and macroevolution^12^, but importantly the same studies^11^ also note that some form of stabilizing selection on wing shape must operate to generate the patterns observed. Across a range of taxa, a related finding that within-species evolvability is a strong predictor of macroevolution can only be fully resolved by invoking stabilizing selection^3^.

**Figure 1.**
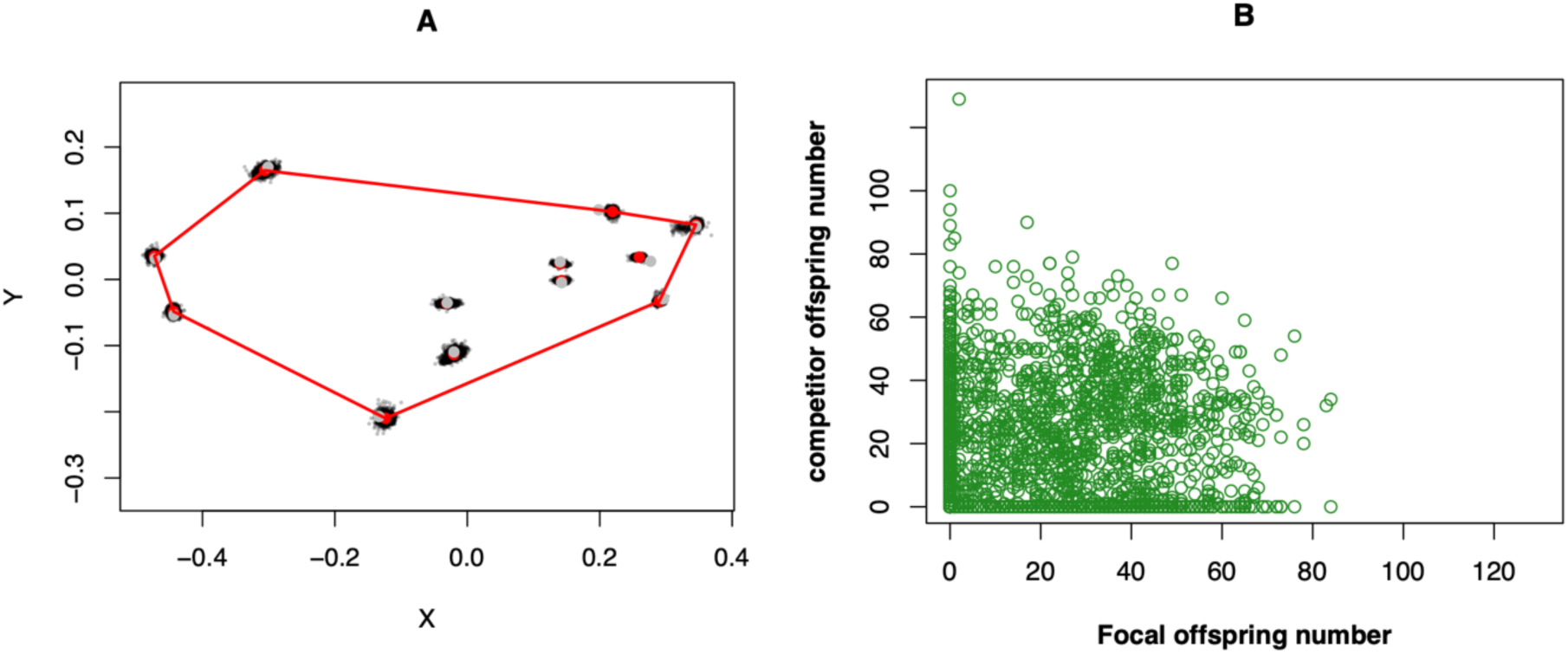
Variation in wing shape and fitness in *D. melanogaster*. Panel A shows wing shape variation represented by 12 landmarks demarking major vein intersections, here scaled and rotated in a Procrustes analysis (N = 1994). Panel B shows focal and competitor offspring number from a competitive mating assay; this is a binomial measure of competitive reproductive success (N = 2130).

We find pervasive stabilizing selection on multivariate wing shape in *D. melanogaster*. Assessing the same landmark coordinates rotated into the same shape space as past work^8,9,11^, as well as competitive lifetime reproductive success in the same individual flies, we find that estimates of quadratic selection differentials are nearly universally negative across dimensions of wing shape and related strongly to the amount of phenotypic variance (Figure 2A). While the pattern is strongest when selection is estimated on phenotypic scores on the phenotypic covariance matrix **P,** we find the same pattern when selection is inferred on phenotypic scores on existing^11^ estimates of the genetic covariance matrix **G**, the mutational covariance matrix **M**, and the macroevolutionary rate matrix **R** (Figure 2A). These results (which are robust to variance-standardization; Extended Data Figure 1) indicate that the amount of stabilizing selection is proportional to the amount of variance in any given combination of wing shape traits, consistent with a model of stabilizing selection against total multivariate distance from the mean wing shape. To asses this conjecture directly, we fit models of selection on total squared Euclidean distance (calculated as the sum of squared principal component scores for each individual). We found significant selection against multivariate distance (model without sex interaction; glmm coefficient= -0.0012196 CI =[-0.0017352, -0.0005996], P_mcmc_ < 0.001) that is sex specific; selection is an order of magnitude stronger in males than females (model with sex interaction; glmm interaction coefficient= -0.0019072, CI = [-0.0029554, -0.0007604], P_mcmc_ < 0.001; Figure 2B). We obtain the same conclusions in an analysis of fecundity, instead of competitive reproductive success, as a fitness measure (Extended Data Figure 2). Together, these results indicate a pattern of stabilizing selection against extreme wing shape, driven primarily by variance in male fitness.

**Figure 2.**
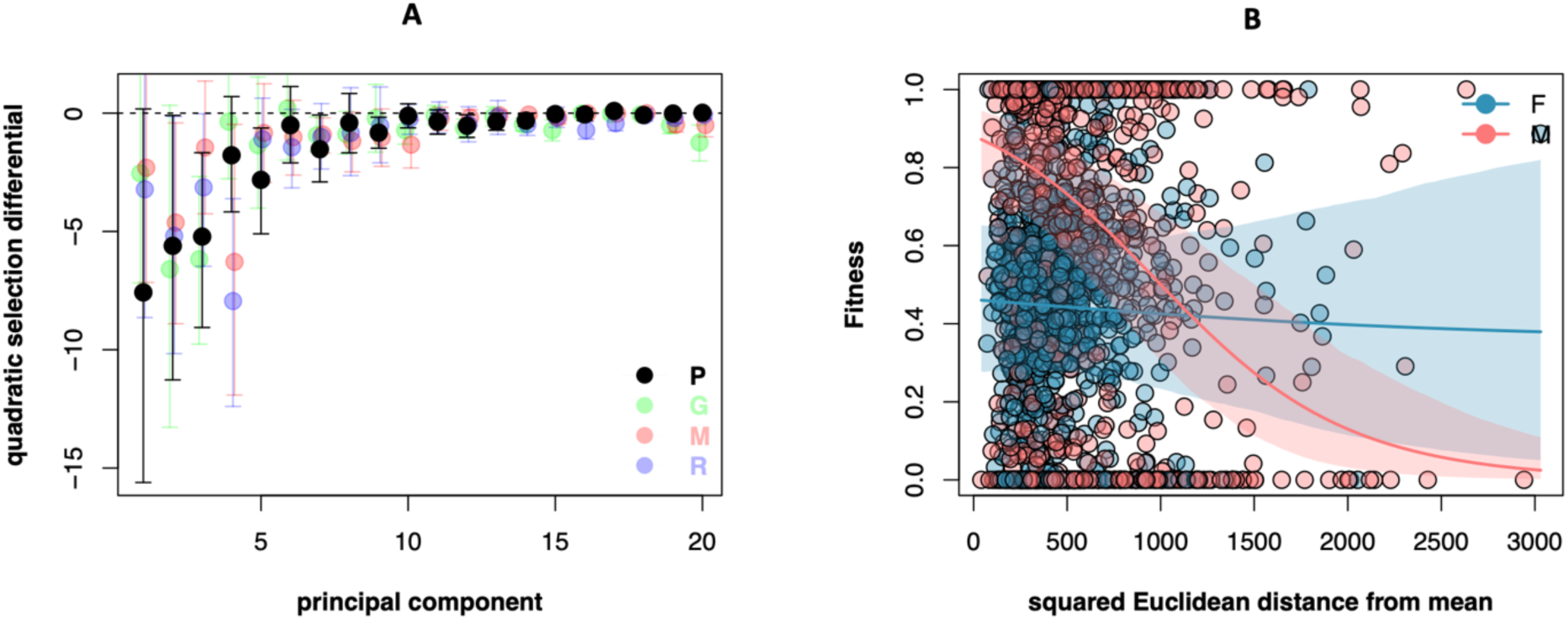
Pervasive stabilizing selection on wing shape is proportional to phenotypic variance. Panel A shows nonlinear selection differentials for raw phenotypic scores on the principal components of the Phenotypic (P), genotypic (G), mutational (M), and macroevolutionary (R) covariance matrices. Panel B shows sex-specific selection against total Euclidean distance from the mean (calculated as the sum of squared scores on the principal components of P).

We propose the stabilizing selection we have observed is likely apparent, and generated by pleiotropic effects of segregating variants on other traits directly affecting fitness^13–15^. Three lines of evidence support this interpretation. First, we have measured selection on wings in a benign lab environment to which the population is well adapted (>500 generations), an environment in which strong direct selection on wing shape is not expected. Although males can use wings to sing and display, interference color display patterns are largely orthogonal to wing shape^16^, and there is unlikely to be selection on flight performance in standard fly vials in which fitness was assayed. Second, we find that stabilizing selection is proportional to standing variance, a pattern consistent with apparent selection via pleiotropy; under such a model of apparent selection, stabilizing selection is predicted to be highest on trait combinations with high variance^13^, as segregating variants affecting these combinations by definition are highly pleiotropic and are most likely to have large pleiotropic effects on other traits that directly affect fitness. Third, we find no detectable selection on major-effect mutations on wing vein structure: in an analysis of 34 flies (14F, 20M) exhibiting cross-veinless phenotypes, we find no reduction in fitness compared to wildtype (N = 1935) vein structure (effect of veinlessness = 0.6689 CI = [-0.7538 2.0325], P_mcmc_ = 0.314 Extended Data Figure 3). This lack of selection on major effect mutations directly affecting wing structure, but detectable stabilizing selection on quantitative wing shape variation, is most consistent with a model of apparent stabilizing selection via pleiotropy.

Such an inference of apparent stabilizing selection in an adapted population represents a background force expected to constrain divergence. We can then assess directly whether our estimate of total stabilizing selection (Lande-Arnold^17^ estimate of γ = -0.00017, 95% CI = - 0.00033 – -0.0000088, not variance-standardized), when combined with inference of standing genetic variance^11^, is generally consistent with patterns of deep-time divergence in wing shape across the Drosophilidae. To do this we explore the simplest model of divergence under stabilizing selection, an Ornstein-Uhelenbeck (OU) process^18^, where lineages diverge under genetic drift proportional to 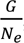, and are pulled back towards the optimum by 𝛼. Here 𝛼 is the curvature of the adaptive landscape, which can be defined^2,6^ by 𝛼 = −𝛾, where 𝛾 is the nonlinear selection gradient^17^ estimated from an individual fitness surface (i.e., our estimate of selection given above). The stationary (i.e., deep-time) variance among lineages under such a process is given as 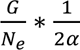, which can be recovered under simulation by rescaling the Drosophilidae phylogeny from millions of years to units of generations (Extended Data Figure 4). Stationary variances under the estimated value of stabilizing selection and **G** are plotted against a range of effective population sizes in Figure 3. We find that our estimate of selection readily explains the constrained divergence in wing shape across the clade; low (but reasonable for many wild *Drosophila* populations^19^) values of *N_e_* produce an expected stationary variance consistent with the observed^11^ among-species phenotypic variance. This analysis can be seen as a proof of concept demonstrating that the observation of low divergence among species coupled with significant standing variance within species can be explained by observable levels of stabilizing selection.

**Figure 3.**
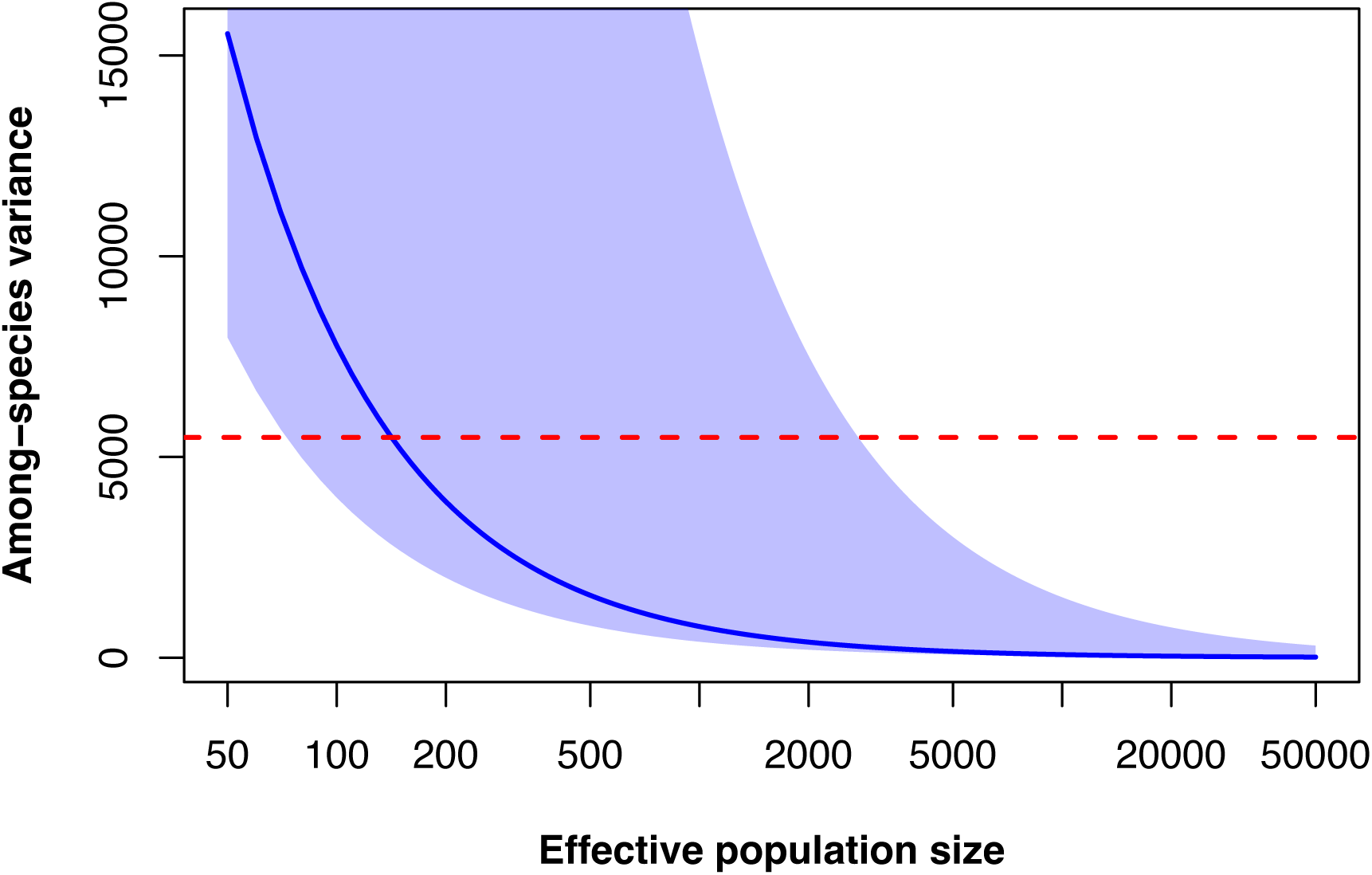
A macroevolutionary model of genetic drift plus stabilizing selection can explain low levels of observed divergence across the clade. The stationary variance (i.e., predicted among-species variance at equilibrium) from an Ornstein-Uhlenbeck model parameterized with the estimate of multivariate stabilizing selection (pooled by sex) as a function of Ne. Shaded area shows the 95% confidence interval of the estimate of selection; dotted red line shows the observed phenotypic variance among species.

Our results reconcile some aspects of fly wing evolution, yet also highlight some lingering challenges. First, our selection estimates cannot explain why some dimensions of wing shape exhibit conserved levels of high variance while others exhibit low variance. While conserved stabilizing selection could explain such a pattern, it would lead to the opposite pattern across principal components of wing shape to that which we observed, with the weakest selection observed to act on the components with the most standing variance^20,21^. It is more likely that aspects of development play a leading role in the conserved eigen structure of wing variance^8,9^. Second, while a simple OU model parameterized with our estimate of total selection can reign in predicted divergence among species to the levels observed in nature, this model is clearly not a complete or accurate description of wing evolution. It seems untenable that all flies have evolved to a single global optimum wing shape, and the short phylogenetic half life (4058 generations, a miniscule fraction of the total depth of the Drosophilid phylogeny) is inconsistent with the observation^11^ of moderate to high phylogenetic signal in wing shape. Finally, *N_e_* on the order of hundreds to ∼2000 individuals is required to predict among species variance congruent with observed vales; while this is consistent with e.g. wild *D. buzzatii*^19^, it is inconsistent with the large *N_e_* typically inferred for *melanogaster.* It is unclear what a reasonable ancestral mean value of *N_e_* would be for the clade. These last three problems could be resolved by a multi- or fluctuating-optimum^3^ OU process, which would lead to increases in predicted among-species variance at a given *N_e_* and also increase phylogenetic signal. However, we lack any first principles for how optima should be placed or move on an adaptive landscape^22^.

Stabilizing selection on wing shape in our experiment was driven almost entirely by males, which also exhibited greater variance in fitness than females (Levene’s test *F*_1,1836_ = 747, *P* < 0.0001). This finding of sex-specific selection explains the discrepancy between our findings of pervasive stabilizing selection, and those of Rohner and Berger^9^, who found no evidence of stabilizing selection on the same wing landmarks in the distantly-related but phenotypically similar *Sepsis punctum*; their analysis focused only on female fitness, rather than competitive male reproductive success. Although typically viewed as a diversifying force^23^, our results demonstrate that sex-specific selection can act as a force constraining the evolution of phenotypic diversity.

## Methods

Flies were *D. melanogaster* of the Larry Harshman Moderate density (LHm) background. This lineage of flies was collected in California in the early 1990s^24^ and has been maintained since the mid-1990s at large, density-controlled population size under 25C temperature and standard *Drosophila* food media. This lineage is lab adapted (over 500 generations in the lab) and harbors substantial genetic and phenotypic variance in most traits, including fitness.

### Fitness assay

We conducted a competitive fitness assay in which a focal, wildtype, unmated, male or female fly was placed in a vial with two other unmated flies, a male and female homozygous for the *bw* eyecolor mutation but otherwise of LHm background. Thus each focal fly was placed with a brown eyed competitor of the same sex, as well as a brown eyed mate of the opposite sex. Following two days of mating interaction, all flies were removed and resulting offspring were allowed to develop, and scored for eye color after eclosion. We thus obtained a binomial measure of reproductive success for each focal fly, although we also present an analysis of raw fecundity (see below). Our measurement of fitness via competitive reproductive success over a two day window is generally consistent with lifetime reproductive success in the context of propagation history of the LHm lineage, where the offspring founding a new generation are obtained after 18 hours of egg laying following two days of mating interaction. The fitness assay was blocked by generation (n = 8) and conducted at room temperature for logistical reasons.

### Wing phenotyping

Following the mating assay focal flies were frozen, and later the right wing (left, if the right was damaged) was removed, dry mounted under a standard cover slip, and photographed under transmitted light using a stereomicroscope (Olympus SZX-10). The microscope was fitted with an objective revolver allowing the objective to be aligned with the photographic optical path, eliminating the distortion otherwise inherent in imaging through a stereomicroscope. A stage micrometer was photographed following the same procedures to scale landmark measurements to units of mm. The 12 vein-intersection fixed landmarks were then placed by hand using *geomorph*^25^ in R. Severely damaged wings or those missing crossveins were excluded, although we did include a small number of wings which had a missing landmark; for these we used a thin plate spline procedure to estimate the missing landmark using *geomorph*. Landmark coordinates were rescaled from units of raw pixels to millimeters, using the mean of measures from sperate images of a stage micrometer imaged identically to the wings. We then used a Procrustes analysis using *Morpho*^26^ in R to scale and rotate the landmark coordinates to obtain standardized shape variables, using Houle et al.’s^11^ *D. melanogaster* consensus as a target in the Procrustes rotation. We multiplied the resulting raw shape variables by 1000 to be consistent with past work^11^. In order to estimate the phenotypic covariance matrix to perform a principal component analysis, we used a Bayesian multi-response mixed effects model with the vector of shape traits as the response, sex of the fly as a fixed effect and block as a g-side random effect; the r-side unstructured covariance matrix gives an estimate of **P**. We projected eigenvectors of **P**, as well as previous estimates (from ^11^) of **G**, **M**, and **R**, on the centered shape data to obtain phenotypic scores on the principal components of these matrices. Note that due to scaling and rotation inherent in the Procrustes analysis, there are 20 non-null dimensions of shape variation each of these covariance matrices.

### Statistical analysis

We estimated unstandardized quadratic selection differentials, conditioned on sex, separately for each principal component by fitting separate generalized mixed effects modes each with sex as a fixed effect, as well as linear and quadratic terms for pc score. Block was included as a random effect. The response variable was the vector of red eyed and brown eyed offspring number. We assumed a binomial distribution for the response variable, and estimated the residual variance (instead of fixing to unity), as the response was not a single Bernoulli trial but rather many for each data point, and so overdispersion is possible (even likely)^16^. We then approximated^16,27^ the nonlinear selection differential^21^ *C* from the quadratic regression coefficient (*c*) as 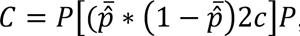, performing this across the posterior distribution to obtain credible intervals. We also report estimates of variance-standardized quadratic selection differentials in Extended Data Figure 1.

We then estimated selection on total Euclidian distance by fitting a binomial glmm with red and brown eyed offspring number as the response, sex as a fixed effect, as well the sum of individual pc scores and the sum of squared pc scores as fixed effects. The sum of raw pc scores is necessary to account for any directional selection (although we found no statistically significant directional selection in this analysis), the sum of squared pc scores is a measure of the squared Euclidean distance from the mean wing shape. We re fit this model to include an interaction with sex and the other fixed effects, to test the hypothesis that selection may be sex-specific. We obtained the same conclusions when fitting a similar model, but with the number of red eyed offspring a Poisson response variable. In order to obtain a Lande-Arnold^17^ estimate of nonlinear selection on the total phenotypic variance for use in an OU model that is most appropriate for approximating the adaptive landscape^28^, we fitted a linear mixed model with the sum of raw pc scores and squared Euclidean distance as fixed effects. We modeled relative fitness as a Gaussian response variable, where relative fitness was calculated as the proportion of Red eyed offspring divided by the mean proportion, calculated separately for males and females^29^. Total offspring number was included as a weight term. We then used two times^30^ the estimate of the nonlinear regression term as our estimate of γ; this gave us a classical Lande-Arnold^17^ estimate of nonlinear selection, pooled by sex, that avoids the challenges^16^ of approximation of nonlinear selection gradients from a glm regression coefficient.

To assess fitness effects of cross vein mutations, we fitted a binomial mixed effects model to an expanded dataset that included flies with missing crossveins. These flies were excluded from the landmark analysis as all of the landmarks could not be placed on these mutant individuals. Our model included wing type (wild type or crossvein mutant) as a fixed effect. We plot the data (Extended Data Figure 3) using the log odds of fitness (ln(red+1 /brown +1)) as this is easier to visualize.

For our assessment of OU model fits, we obtained an estimate of total within-species genetic variance and among-species phenotypic variance as the sums of the diagonals of the **G** and **R** (respectively) matrices for wing shape obtained from ref ^11^, as well as our own estimate of 𝛼 (see above). We confirmed the stationary expectation for the OU process by simulating 1000 runs of OU evolution assuming an *N_e_* of 142, calculated as the *N_e_* required to produce a theoretical stationary variance equivalent to the observed among-species variance (=5487) given our estimate of selection. Simulations (Extended Data Figure 4) were performed in *phytools*^31^, using the a time-calibrated Drosophilid phylogeny obtained (by vcv2phylo() in *ape*^32^) from the phylogenetic covariance matrix presented in ref ^11^. We rescaled the branch lengths from millions of years to generations assuming 10 generations per year for wild *Drosophila*.

All analyses were performed in R. Mixed models were fitted by Bayesian MCMC in *MCMCglmm*^33^, with the exception of the linear mixed model used to estimate γ which was fitted by REML using *lme4*^34^. Large language models (ChatGPT) were used as a coding aid in modifying R code for producing visually appealing figures for some of the figures.

## Acknowledgements

We thank David Berger, Masahito Tsuboi, and John Stinchcombe for discussion and statistical advice. Funding was provided by grants from the Swedish Research Council (Vetanskapsrådet: grants no. 2019-03706 and 2024-04599), the Crafoord Foundation (grant no. 20220602), and KK-stiftelsen (grant no. 2024STG)

## Author contributions

A.B. and S.P.D. designed the study. A.B. performed the experiment, collected and processed the data. S.P.D. analyzed the data and drafted the paper. Both authors revised the paper.

## Extended Data Figures

**Extended Data Figure 1.**
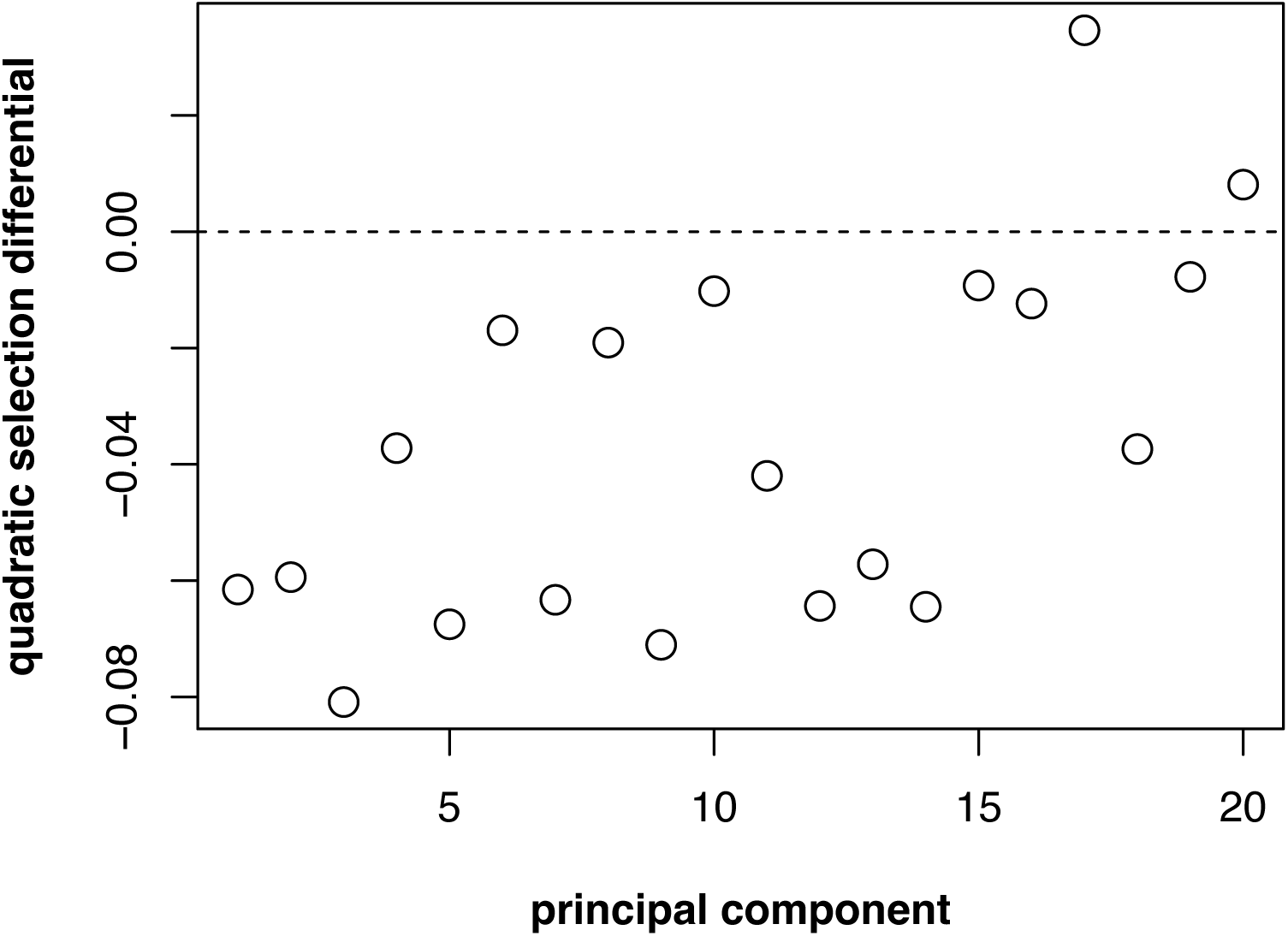
Results are robust to variance standardization. Point estimates of nonlinear selection differentials for phenotypic scores on the principal components of the P matrix, where the scores for each pc have been variance standardized prior to estimation of the selection differentials.

**Extended Data Figure 2.**
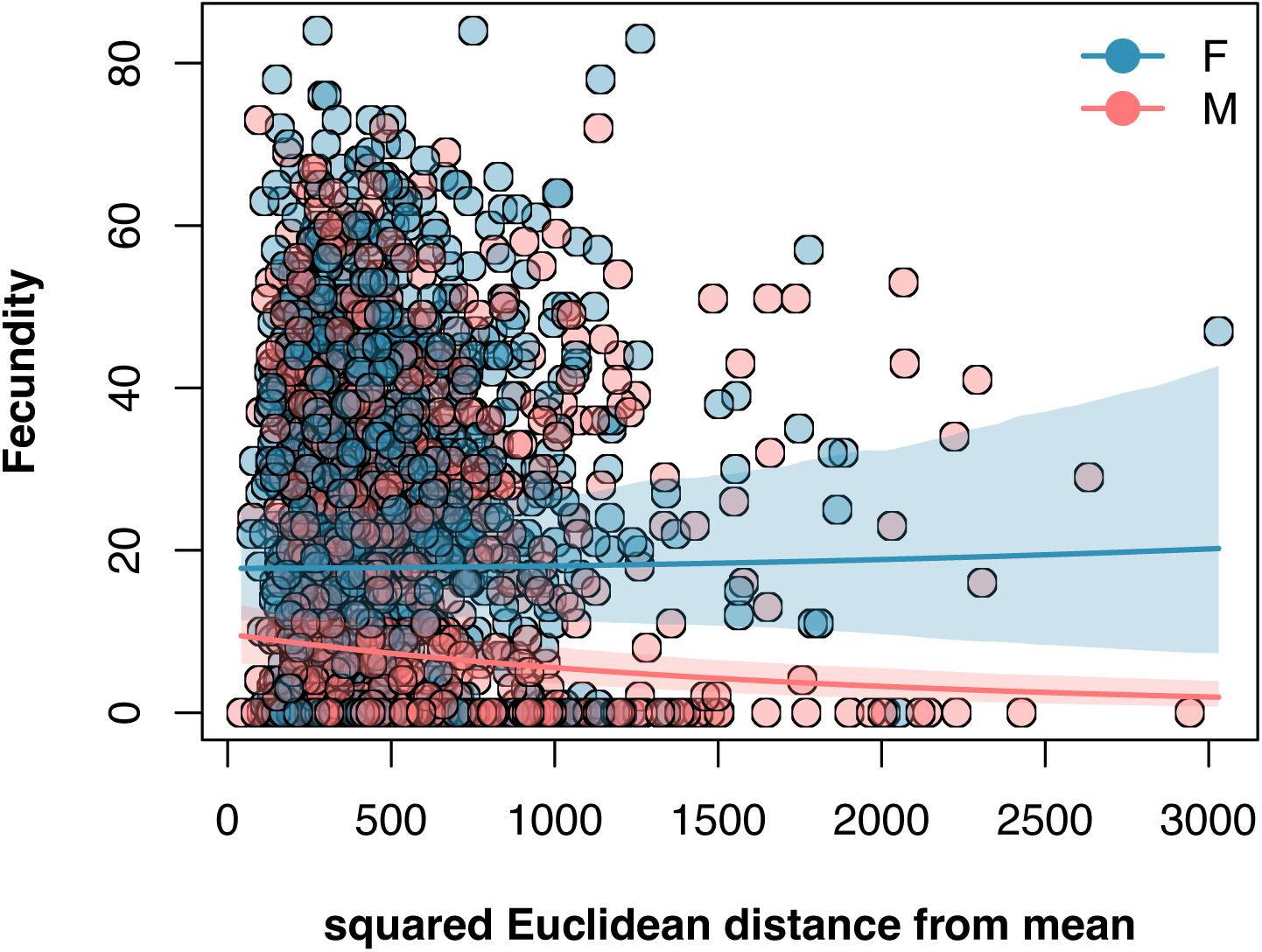
Sex-specific selection against phenotypic extremes, assuming raw fecundity as a fitness measure. Shown are the model predicted values and 95% credible intervals from a Poisson generalized mixed model, with otherwise-identical structure that that fitted for Figure 2b.

**Extended Data Figure 3.**
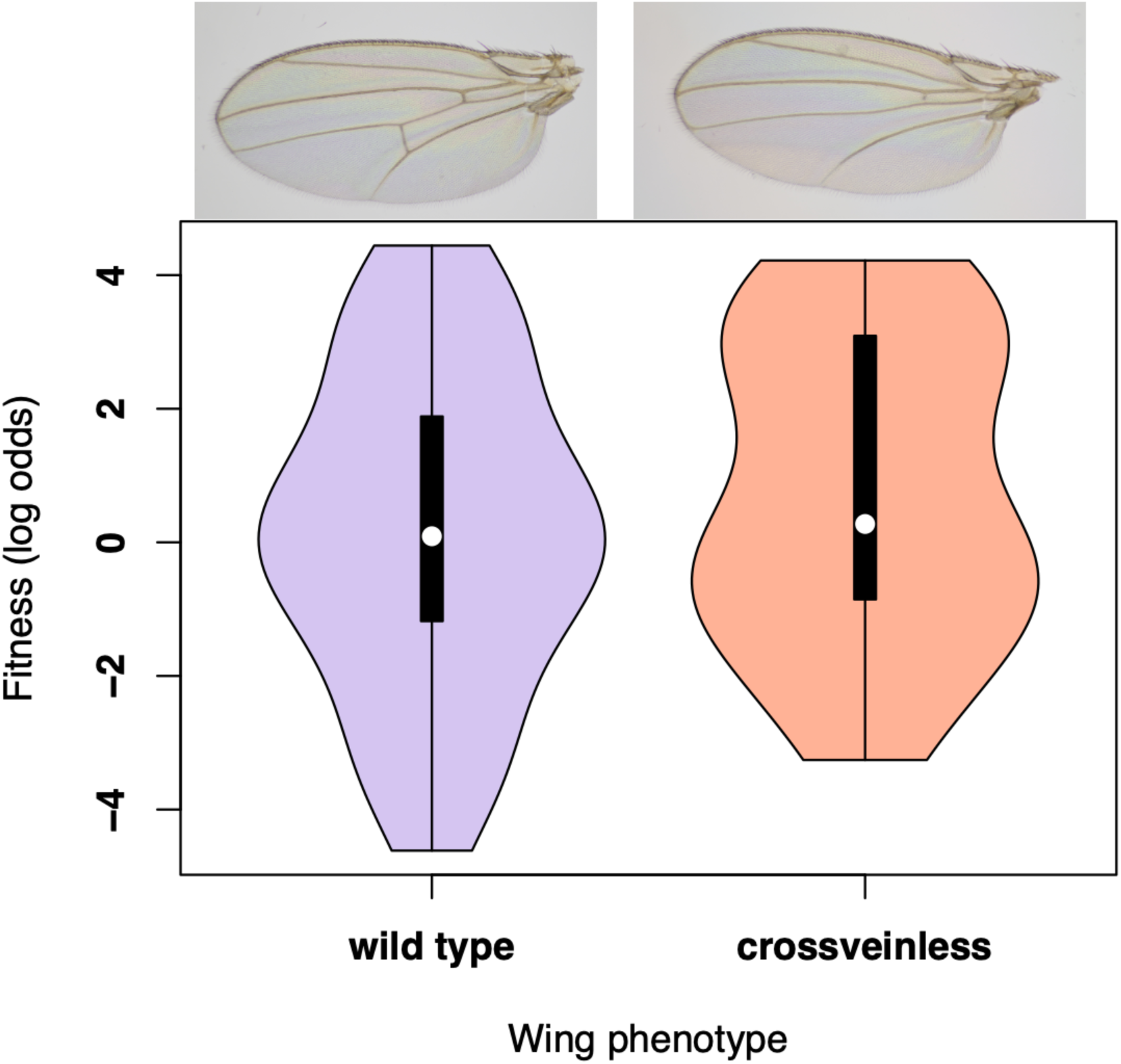
No selection against missing crossveins. Shown are the log odds of fitness for flies with wild type and crossveinless wings. A binomial generalized mixed effects models revealed no evidence of selection against the cross veinless phenotype.

**Extended Data Figure 4.**
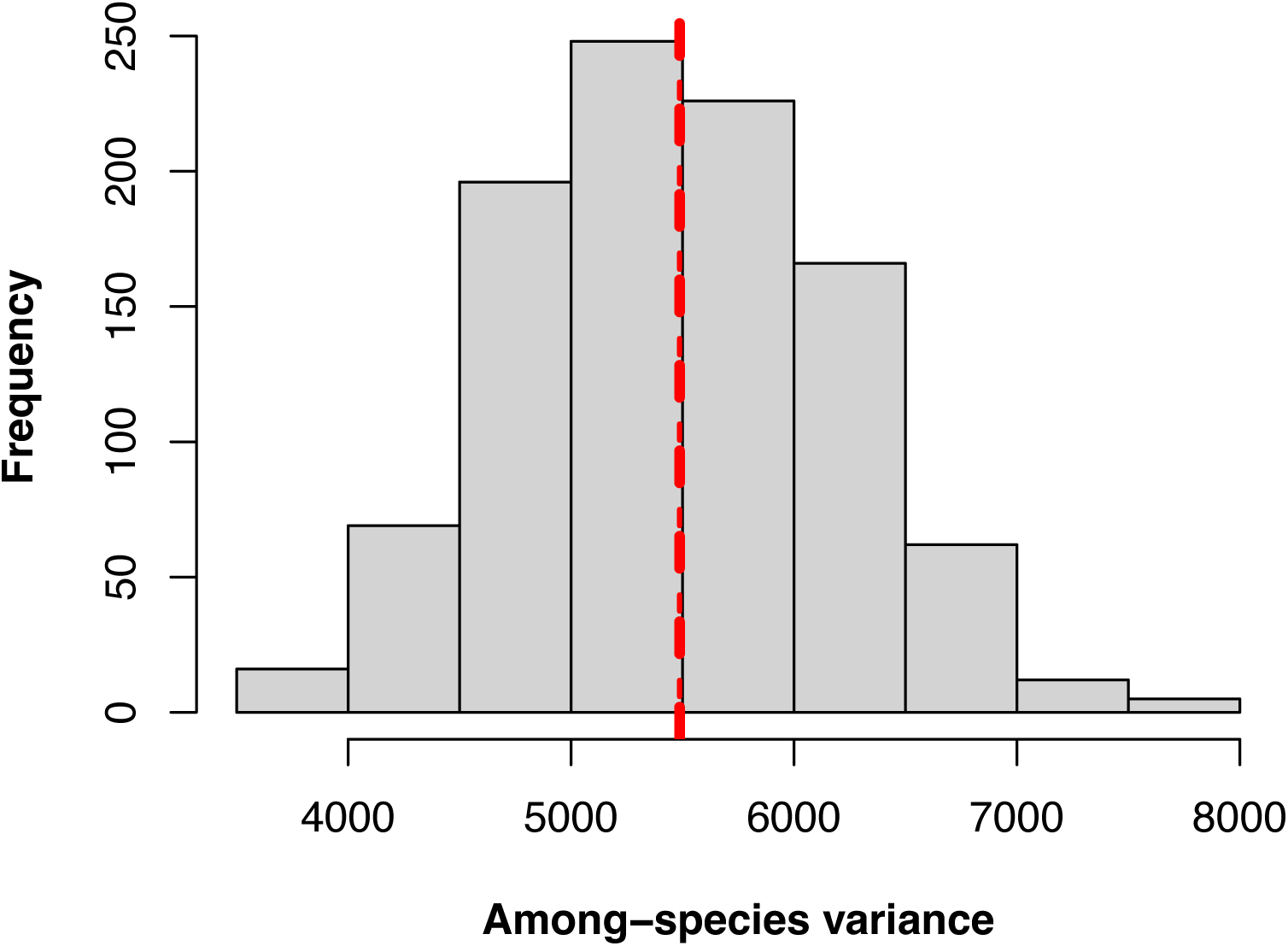
Correspondence between theoretical stationary OU variance and simulated OU evolution on the Drosophilid phylogeny. Red line shows observed among-species phenotypic variance. Histogram shows frequency distribution of simulated (1000x) OU evolution on the rescaled Drosophilid phylogeny assuming calculated from the intersection of the two lines in Figure 3 (see Methods).

## Notes

### Competing Interest Statement

The authors have declared no competing interest.

